# Calcitonin Gene-Related Peptide (CGRP) Receptor Antagonism Reduces Motion Sickness Indicators in Mouse Migraine Models

**DOI:** 10.1101/2022.06.03.494762

**Authors:** Shafaqat M. Rahman, Anne E. Luebke

## Abstract

**Background:** Migraine and especially vestibular migraine are associated with enhanced motion sickness and motion-induced nausea. Murine models of migraine have used injections of Calcitonin Gene-Related Peptide (CGRP) or other migraine triggers [i.e., sodium nitroprusside (SNP)] to induce migraine sensitivities to touch and light. Yet, there is limited research on whether these triggers increase motion-induced nausea, and if migraine blockers can reduce these migraine symptoms. We hypothesized that systemic delivery of CGRP or SNP will increase motion sickness susceptibility and motion-induced nausea in mouse models, and that migraine blockers can block these changes induced by systemically delivered CGRP or SNP.

**Methods:** We investigated two measures of motion sickness assessment [motion sickness index (MSI) scoring and motion-induced thermoregulation] after intraperitoneal injections of either CGRP or SNP in C57B6/J mice. The drugs olcegepant, sumatriptan, and rizatriptan were used to assess the efficacy of migraine blockers.

**Results:** MSI measures were confounded by CGRP’s effect on gastric distress. However, analysis of tail vasodilations as a surrogate for motion-induced nausea was robust for <u>both</u> migraine triggers. Only olcegepant treatment rescued tail vasodilations.

**Conclusion:** These preclinical findings support the use of small molecule CGRP receptor antagonists for the treatment of motion-induced nausea of migraine, and show that triptan therapeutics are ineffective against motion-induced nausea of migraine.

## Introduction

Motion sickness and motion-induced nausea are symptoms of migraine and especially vestibular migraine (VM) ^1–7^. VM patients show lower motion perception thresholds compared to healthy controls and also show enhanced susceptibility to motion sickness^3^. Interestingly, Wang and Lewis (2016) found that in VM patients but not in migraine or healthy controls, the residual sensory conflict between gravitational (otolith) and rotational (semicircular canal) cues correlated with motion sickness susceptibility ^8^.

The diagnostic criteria for human motion sickness have recently been updated ^9^ and are included in the international classification of vestibular disorders. The major symptoms of motion sickness include facial pallor, nausea, vomiting, gastric awareness and discomfort, sweating, and hypothermia. Motion sickness-related hypothermia is broadly expressed phylogenetically in humans, mice, rats, and musk shrews ^10, 11^. However, rodent models of motion sickness have been constrained because rodents do not have an emesis reflex. Pica-eating behavior is thought to be an alternative to vomiting, yet, pica has not been shown to be a sensitive measure of motion sickness ^12^. Instead, piloerection, tremor, fecal, and urinal incontinence have been used in scoring criteria called the motion sickness index (MSI) to quantify the degree of motion sickness-like behavior experienced by rats and mice to emetic stimuli ^12, 13^. In addition, thermoregulatory changes can also be used to assess motion sickness, which in the mouse model involves a decrease in head temperature (hypothermia) and transient tail-skin vasodilation early in the onset of provocative motion ^11^. Tail vasodilations to provocative motion have been reported to precede emesis events in the musk shrew, serving as an early index for motion-induced nausea in experimental rodent models.

Despite the high prevalence of motion-induced nausea in VM, the underlying mechanisms have yet to be defined. It is accepted that migraine involves the neuropeptide Calcitonin Gene-Related Peptide (CGRP). CGRP is upregulated during migraine attacks^14, 15^, infusion of CGRP can induce migraine ^15^, and antibodies that block CGRP or its receptor can effectively treat most migraines ^16, 17^. Yet, it is unclear if CGRP signaling antagonism can also alleviate motion-induced nausea of VM.

Animal models of migraine have used injections of CGRP or other migraine triggers such as sodium nitroprusside (SNP) - which generates nitric oxide and stimulates the release of CGRP – to induce allodynic responses to touch ^18, 19^ and light-aversive behavior ^20, 21^. However, it is not known if these triggers can induce motion-induced nausea in preclinical models. It is also not known if migraine blockers used in preclinical models of touch and light sensitivities can also block motion-induced nausea and motion sensitivity characterized in VM.

To better understand CGRP’s role in VM motion-sickness susceptibility, we investigated two measures of motion sickness assessment (MSI scoring and motion-induced thermoregulation) after systemic injections of either CGRP or SNP in the wild-type C57B6/J mouse. In this study, we found that MSI measures based on gastric distress were lessened by CGRP antagonism. This was driven by CGRP’s effect in causing diarrhea, which was not observed after SNP administration. However, motion-induced thermoregulation was a robust model of motion sickness. CGRP receptor antagonism by olcegepant was effective in relieving CGRP effects on motion-induced nausea, whereas triptan therapies were not efficacious. Our studies provide a strong premise that antagonizing CGRP signaling will be effective for treating motion-induced nausea of VM, as it has been shown to be highly effective for typical migraine.

## Materials & Methods

### Animals

C57B6/J mice were obtained from Jackson Laboratories (JAX #664) and were housed under a 12-hour day/night cycle under the care of the University Committee on Animal Resources (UCAR) at the University of Rochester. Mice are housed with *ad libitum* access to food, water, bedding, and enrichment materials. A total of 145 mice (73 M/ 72 F) were tested, and all studies were powered sufficiently to detect male/female differences. Prior to behavioral testing, mice were equilibrated in the testing room controlled for an ambient temperature between 22-23°C for at least 30 minutes. Mice were tested around 2.3 to 6 months of age. This age range correlates to 20-30 years in humans and is within the range that migraine symptoms most likely occur in patients ^22^. Different cohorts of mice were used to test motion sickness indices and motion-induced nausea. Testing occurred from 9:00 am to 4:30 pm during the day cycle to control for behavioral and thermoregulatory changes that may arise from circadian rhythms. For motion-induced nausea testing, mice were screened for instances of patchy fur (alopecia) and were not included in this study ^23^.

### Drug administration

All injections were performed intraperitoneally (IP) with a fine 33-gauge insulin syringe. Dulbecco PBS served as the diluent and as the vehicle control. Migraine blockers used were the CGRP-receptor antagonist olcegepant and the selective serotonin receptor agonists sumatriptan and rizatriptan. The concentrations are listed: 0.1x, 0.5x, and 1.0x CGRP were prepared at 0.01, 0.05, and 0.1 mg/kg (rat ɑ-CGRP, Sigma), 1.0x olcegepant was prepared at 1.0 mg/kg (BIBN4096, Tocris), 1.0x sumatriptan was prepared at 0.6 mg/kg (Sigma-Aldrich), 1.0x rizatriptan was prepared at 0.6 mg/kg (Sigma-Aldrich), and 0.1x, 0.5x, and 1.0x sodium nitroprusside (SNP)-(Sigma-Aldrich) were prepared at 0.25 mg/kg, 1.25, and 2.5 mg/kg. After injection, animals were placed in a separate cage from their home cage to recuperate from injection stress. Mice were tested approximately twenty to thirty minutes after delivery of either vehicle, CGRP, SNP, CGRP or SNP co-administered with a blocker, or drug controls. Animals were gently handled and anesthesia was not needed. All animal procedures were approved by the University of Rochester’s (IACUC) and performed in accordance with the standards set by the NIH.

### Off-Vertical Axis Rotation (OVAR)

Prior human and rodent studies have used OVAR to assess the otolith-ocular reflex and assess the semicircular canal-otolith interaction ^24–26^. Constant velocity OVAR at a tilt can be disorienting and promote motion sickness in human participants ^27, 28^, and further studies in mice have shown that provocative rotation leads to pica and observations of urination, piloerection, and tremor ^29^. In this study, a two-cage rotator (cage dimensions: 12.7 cm x 8.9 cm) was use for twenty minutes (60 rpm, 45° tilt from the vertical, 20 cm from the axis of rotation) as an OVAR vestibular challenge.

### Motion-Sickness Index (MSI) Testing

We adapted Yu et al.’s protocol that previously assessed the drugs scopolamine and modafinil on mitigating motion sickness indicators in rats and mice ^12^. We incorporated OVAR (described above) as the vestibular challenge (VC) in this test. We evaluated MSI in mice at the end of the following time points: A) a five-minute baseline (*pre-injection*), B) twenty to thirty minutes after drug injection (*post-injection*), and C) five minutes after the OVAR (*post-VC*). Mice were placed in a testing box separate from their home cage to observe for indicators of motion sickness detailed in **Fig. 1A**. A motion sickness index (MSI) score was determined by the summation of the indicators.

**Fig. 1:**
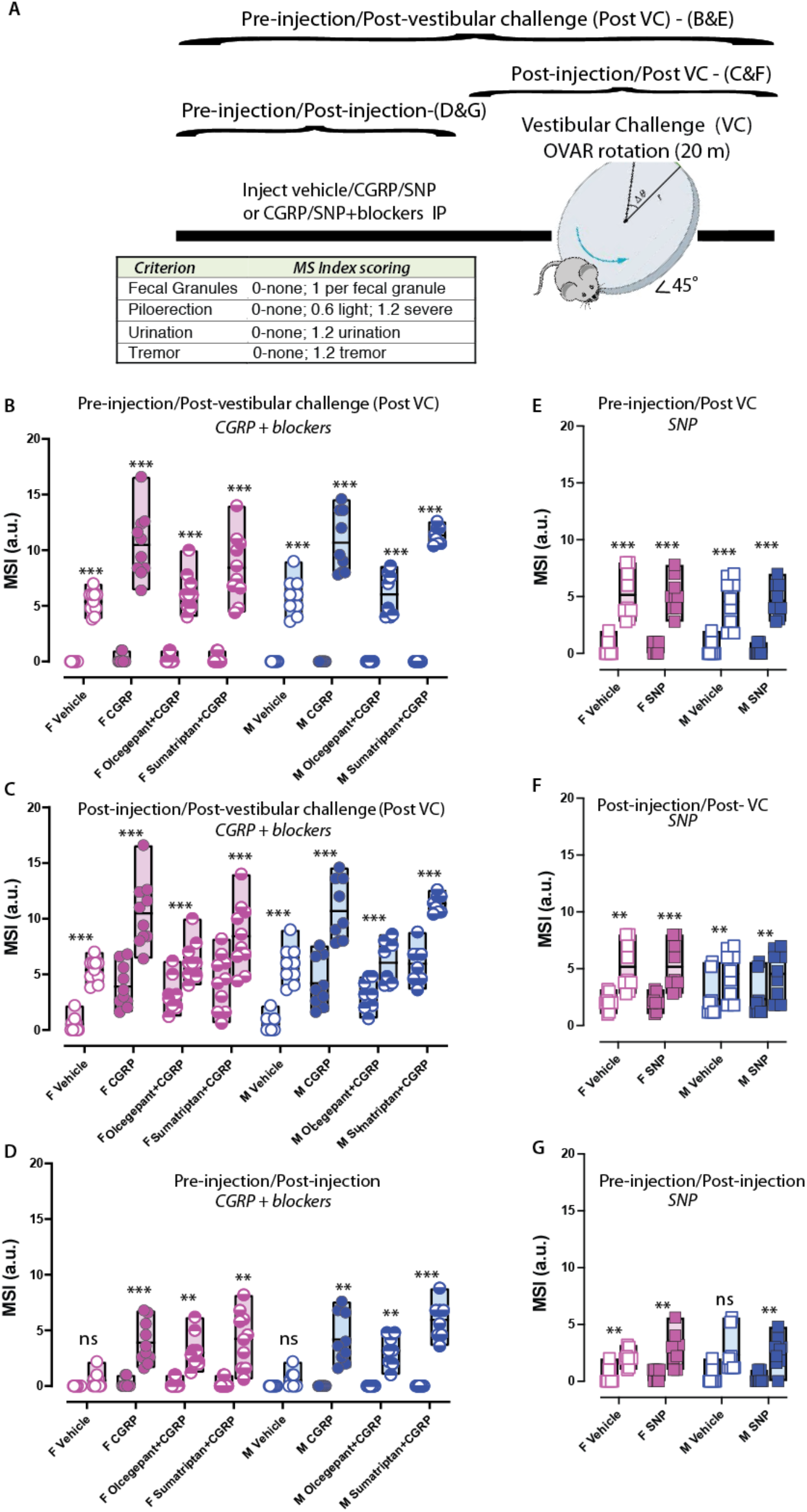
CGRP and SNP’s effects on MSI: Female (pink) and male (blue) C57B6/J mice were tested for motion sickness indicators. Treatments were delivered intraperitoneally (IP): vehicle control (open circle/open square), CGRP (closed circle), SNP (closed square), olcegepant + CGRP (top half-filled circle), and sumatriptan + CGRP (bottom half-filled circle). **(A)** Assay timeline is illustrated and a table of criteria and MSI scoring. Motion sickness index (MSI) was computed by a summation of criterion at the appropriate scoring and involved measuring feces, piloerection, urination, and tremor. MSI was recorded at three different time points. The vestibular challenge involved off-vertical axis rotation (60 rpm, 45° tilt from the vertical) for 30 minutes. In each instance, the first bar graph in each condition is MSI calculated either pre-injection **(B & E),** post-injection **(C & F)**, or pre-injection **(D & G)**, and the second part of bar graphs are either post-vestibular challenge **(B, C, E & F)**, or post-injection **(D & G)**. Two-way ANOVAs were analyzed in males and in females. F-statistics and p-values can be found in **Table 1**.

Certain actions were taken to normalize the weight distribution of MS indicators during reporting. Fecal granules (Fg) were counted separately at each time point. The unit weight per Fg after vehicle injection was used to back-calculate the granule number when the weight of fecal incontinence (in grams) was instead measured, for example, in cases of gastric distress. This calculation was necessary during CGRP testing as mice experienced diarrhea, which made it difficult to count fecal incontinence.

Urination was assigned a 1.2 score and was only counted once throughout the time points. Piloerection was measured as either mild (0.6) or severe (1.2), and tremors were assigned a 1.2.

### Motion-Induced Thermoregulation Testing

We adapted Tu et al.’s protocol who first noticed these thermoregulatory changes when measuring the temperatures of the heads, bodies, and tails of mice to provocative motion ^11^. In this study, head and tail temperatures of C57B6/J mice were measured for a total 45 minutes using a FLIR E60 IR camera (model: E64501). This camera is connected to a tripod and is positioned approximately 43 cm above an open, plexiglass box (mouse box) used to house an individual mouse during testing. Both the tripod and mouse box are securely attached to the shaker’s base. Briefly, baseline measurements were recorded for five minutes prior to the provocative motion (-5 to 0 mins). The provocative motion was an orbital rotation (75 rpm, 2-cm orbital displacement), and mice were recorded for 20 minutes (0 to 20 mins). After 20 minutes, the provocative motion was turned off, and mice were recorded for an additional 20 minutes to measure recovery to baseline (20 to 40 mins) as schematized in **Fig. 2**. Head and tail temperatures were measured after data retrieval using the software FLIR Tools+. Tail and head temperatures were measured within predefined field of views: square region (3x3 mm) for tail, and circular region (10x10 mm) for head. Tail measurements occurred 2 cm from the base of the tail and head measurements occurred at the center of the head image, in between the mouse’s ears. Infrared imaging data was collected every minute during baseline measurements, and every 2 minutes during and after the provocative motion. We quantified thermoregulatory changes to provocative motion by comparing changes in tail vasodilations, and we approximated the magnitude of the head hypothermia based on second order curve fit estimates. Transient increases in the tail temperature of the mouse to provocative motion are referred to as *Δ tail vasodilations* (°C), and were computed by subtracting the baseline tail temperature at time t = 0 minutes (rotation ON) from the max tail temperature measured during the first 10 mins of the rotation (0 ≤ t ≤ 10). In order to facilitate quantification and comparisons of the Δ tail vasodilations between treatment groups, a threshold of 1.5°C was imposed upon the data to make it a binary outcome measure. Tail temperature changes equal to or greater than +1.5°C were designated a Δ tail vasodilation and those less than +1.5°C did not meet the criteria. Mice were excluded from further testing if their Δ tail vasodilation during the vehicle control test did not meet the criteria.

**Fig. 2:**
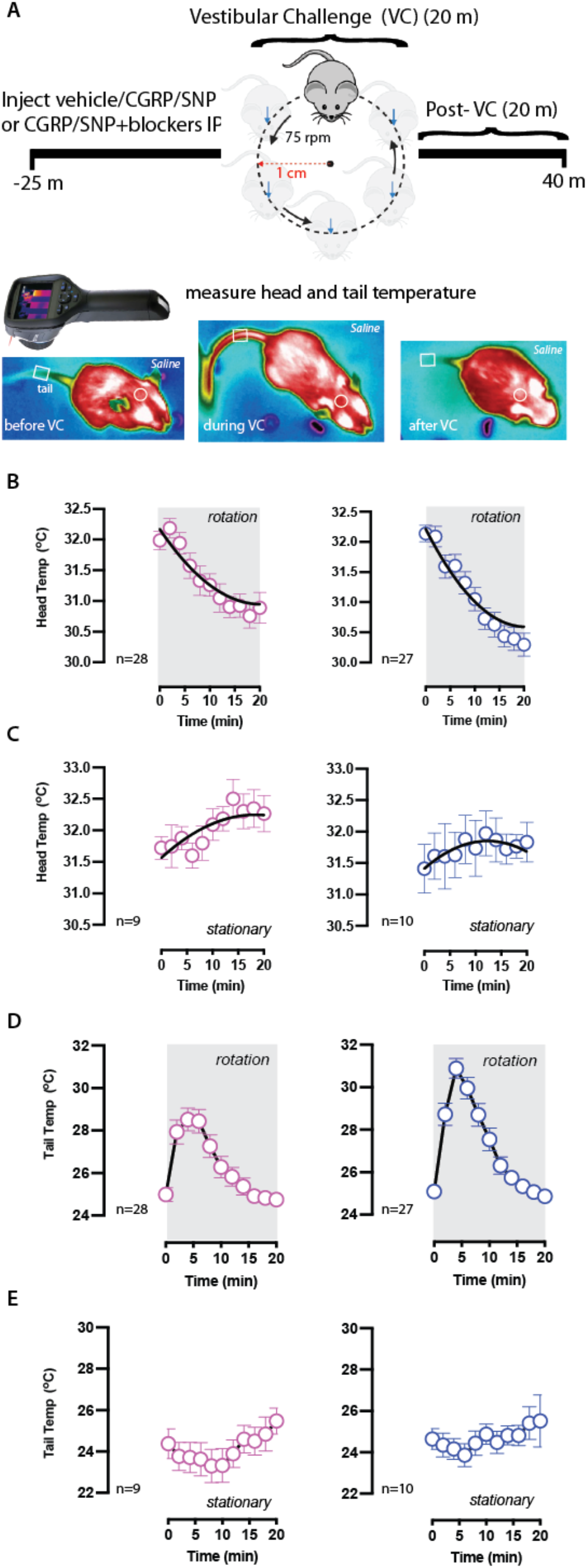
Stationary and provocative motion testing after IP vehicle control: **(A)** Using a FLIR E60 infrared camera, head and tail temperatures for all conditions are recorded before, during, and after a provocative twenty-minute orbital rotation acting as the vestibular challenge. Infrared recordings are 45 minutes in duration. Sample sizes are depicted in the bottom left for female (pink) and male (blue) C57B6/J mice. After administering vehicle (saline), **(B and C)** head and **(D and E)** tail temperatures were compared during **(C and E)** stationary testing or **(B and D)** during the provocative rotation. Head temperatures for stationery and VC testing were fit to the second-order curve (B_2_X^2^ + B_1_X + B_0_) and Δ head temperatures were computed by subtracting temperature at time t = 20 mins from t = 0 mins. Head curves compute similar hypothermia between sexes but different from stationary tests. During the first 10 minutes of the provocative rotation, Δ tail vasodilations were computed to be 4.34 ±0.34°C for females and 5.44 ± 0.42°C for males. Mice during stationary testing did not experience Δ tail vasodilations, and had Δ tail changes of -1.1 ± 0.8°C for females and 0.2 ± 0.6°C for males.

Prior to assessing the efficacy of migraine blockers on thermoregulatory changes due to provocative motion, control experiments were performed to assess i) test-retest reliability of motion-induced thermoregulation and ii) stationary testing (no rotation) after vehicle, 1.0x CGRP, and 1.0x SNP injections. Additionally, we tested motion-induced thermoregulation at 0.1x and 0.5x concentrations of CGRP and SNP to compare with 1.0x effects.

### Statistical Analyses

Statistical analyses were conducted in GraphPad Prism 9.5. Multivariate ANOVAs were analyzed for MSI and motion-induced thermoregulation testing and are described in detail in the results. Second-order curve fitting (B_2_*X^2^ + B_1_*X + B_0_) were used to generalize trends in head temperature profiles and approximate the magnitude of head hypothermia, a single value with no standard error, computed by subtracting the curve’s predicted head temperature at t = 20 mins from t= 0 mins. Head recovery (mins) was approximated by calculating the time to reach baseline using the equation (-B_1_ + √(B_1_^2^)) / (2B_2_) and subtracting it from t = 20 mins. For test-retest reliability of head and tail profiles, Pearson’s correlation coefficient is listed as r (df), where r is the coefficient and df is the degrees of freedom. R^2^ values are provided. Values are reported as mean ± SEM unless noted otherwise, and significance was set at p < 0.05 for all analyses.

## Results

### CGRP and SNP’s effects on motion sickness index

We studied the effects of CGRP release (n = 9M/10F) on motion sickness indicators (MSI) and whether the CGRP receptor antagonist olcegepant or the 5HT_1D_ receptor agonist sumatriptan could mitigate MSI according to the timeline shown in **Fig. 1A**. In females, a two-way ANOVA analyzed MSI outcomes between treatments (CGRP versus CGRP co-delivered with blockers) and between pre-injection versus post-vestibular challenge (VC) time points (**Fig. 1B**). Significant effects were observed due to treatment (F (2.15, 19.35) = 9.94; p < 0.001) and post-VC (F (1.00, 9.00) = 308.20; p < 0.001). Bonferroni post-hoc test showed that this increase was present for every treatment (p < 0.001 for all comparisons). Post-VC MSI was higher after 1.0x CGRP (p < 0.002) and after 1.0x CGRP + sumatriptan (p = 0.1, trending) than vehicle, but significant differences were not observed between 1.0x vehicle and 1.0x CGRP + 1.0x olcegepant. This analysis was also conducted in males and the factors of treatment (F (3.00, 30.00) = 19.09; p < 0.001) and post-VC effects (F (1.00, 30.00) = 634.8; p < 0.001) were also strong. Bonferroni’s post-hoc test showed post-VC MSI was higher than pre-injection MSI for every treatment in males (p < 0.001). Like females, post-VC MSI after 1.0x CGRP and 1.0x CGRP + 1.0x sumatriptan in males was significantly higher than after vehicle (p < 0.001 each), but this difference to vehicle was not seen with 1.0x CGRP + 1.0x olcegepant. In **Fig. 1C**, MSI outcomes after IP injection were compared to post-VC MSI, and 2-way ANOVAs in females and males showed similar trends as was established in the analysis portrayed in **Fig. 1B**. In **Fig. 1D**, pre-injection MSI was compared to post-injection MSI, and with the exception of vehicle, post-injection MSI was higher than pre-injection baseline. **(Fig. 1E-G)** A separate group of mice were tested for MSI after 1.0x SNP (n = 9M/9F), and significant differences were observed when comparing pre-injection to post-VC, post-injection to post-VC, and pre-injection to post-injection. However, the effect of 1.0x SNP was not different from vehicle, and so MSI was not further evaluated with co-delivery of SNP and blockers. A detailed breakdown of F-values and p-statistics is found in **Table 1**.

**Table 1:** ANOVA analyses are depicted for motion sickness index (MSI) testing (top half) and for Δ tail vasodilations from motion-induced thermoregulation assay (bottom half). Analyses were conducted separately in males and females. The following abbreviations are listed: pre-injection is pre-inj, post-injection is post-inj, post-vestibular challenge is post-VC, olcegepant is olceg, and sumatriptan is suma. For MSI, two-way repeated measure ANOVAs with Bonferroni post hoc comparisons were computed to assess two factors: treatment (vehicle, CGRP, CGRP+olceg, CGRP+suma) and timepoint differences (pre-inj vs post-VC, post-inj vs post-VC, and pre-inj vs post-inj). For Δ tail vasodilations, 1-way ANOVAs were conducted to study effects of olcegepant, sumatriptan, and rizatriptan. Dunnett post-hoc comparisons were between treatments to vehicle. F and p-values are listed accordingly.

| Assay | Biological Sex | ANOVAs | Post-Hoc |
| --- | --- | --- | --- |
| Motion Sickness Index (MSI) | Females | pre-inj vs post-VC: $F(1.00, 9.00) = 308.2, p < 0.001$ | vehicle <0.001<br>CGRP <0.001 |
| | | treatment: $F(2.15, 19.35) = 9.94, p < 0.001$ | CGRP + Olceg <0.001<br>CGRP + Suma <0.001 |
| | | post-inj vs post-VC: $F(1.00, 9.00) = 296.0, p < 0.001$ | vehicle <0.001<br>CGRP <0.001 |
| | | treatment: $F(2.10, 19.00) = 10.0, p < 0.001$ | CGRP + Olceg <0.001<br>CGRP + Suma <0.001 |
| | | pre-inj vs post-inj: $F(1.00, 9.00) = 91.0, p < 0.001$ | vehicle 0.200<br>CGRP <0.001 |
| | | treatment: $F(2.00, 18.00) = 9.0, p = 0.002$ | CGRP + Olceg 0.001<br>CGRP + Suma 0.001 |
| | Males | pre-inj vs post-VC: $F(1.00, 30.00) = 634.8, p < 0.001$ | vehicle <0.001<br>CGRP <0.001 |
| | | treatment: $F(3.00, 30.00) = 19.1, p < 0.001$ | CGRP + Olceg <0.001<br>CGRP + Suma <0.001 |
| | | post-inj vs post-VC: $F(1.00, 8.00) = 458.0, p < 0.001$ | vehicle <0.001<br>CGRP <0.001 |
| | | treatment: $F(2.00, 16.00) = 20.0, p < 0.001$ | CGRP + Olceg <0.001<br>CGRP + Suma <0.001 |
| | | pre-inj vs post-inj: $F(1.00, 60.00) = 157.0, p < 0.001$ | vehicle 0.280<br>CGRP 0.002 |
| | | treatment: $F(2.20, 45.00) = 17.0, p < 0.001$ | CGRP + Olceg 0.001<br>CGRP + Suma <0.001 |
| $\Delta$ Tail Vasodilations | Females (CGRP) | Olcegepant: $F(1.21, 10.90) = 10.0, p = 0.007$ | veh vs CGRP <0.0001<br>veh vs CGRP + Olceg 0.057 |
| | | Sumatriptan: $F(1.84, 12.90) = 40.4, p < 0.0001$ | veh vs CGRP <0.001<br>veh vs CGRP + Suma <0.001 |
| | | Rizatriptan: $F(1.23, 7.36) = 24.4, p = 0.001$ | veh vs CGRP 0.005<br>veh vs CGRP + Riza 0.003 |
| | Females (SNP) | Olcegepant: $F(1.31, 9.19) = 7.7, p = 0.016$ | veh vs SNP 0.009<br>veh vs SNP + Olceg 0.208 |
| | | Sumatriptan: $F(1.87, 13.10) = 7.0, p = 0.009$ | veh vs SNP 0.009<br>veh vs SNP + Suma 0.332 |
| | Male (CGRP) | Olcegepant: $F(1.18, 10.60) = 7.6, p = 0.016$ | veh vs CGRP 0.001<br>veh vs CGRP + Olceg 0.315 |
| | | Sumatriptan: $F(1.67, 15.00) = 20.8, p < 0.0001$ | veh vs CGRP 0.000<br>veh vs CGRP + Suma 0.004 |
| | | Rizatriptan: $F(2.00, 11.90) = 25.4, p < 0.0001$ | veh vs CGRP 0.001<br>veh vs CGRP + Riza 0.003 |
| | Male (SNP) | Olcegepant: $F(1.10, 7.67) = 3.6, p = 0.094$ | veh vs SNP <0.001<br>veh vs SNP + Olceg 0.472 |
| | | Sumatriptan: $F(1.06, 7.41) = 2.7, p = 0.143$ | veh vs SNP <0.001<br>veh vs SNP + Suma 0.903 |

### Motion sickness (M SI) outcome is driven by feces criteria

A closer observation of MSI outcomes showed that the main driver of MSI is the magnitude of fecal incontinence, and so 2-way ANOVAs were used to assess differences in fecal incontinence after treatment and across time points in males and females (data not shown). Consistent in both sexes, Bonferroni post-hoc analyses comparing post-VC fecal incontinence outcomes indicated high fecal excretion after 1.0x CGRP (p < 0.001) and after 1.0x CGRP + 1.0x sumatriptan (p < 0.001) compared to vehicle tests, but not in mice treated with 1.0x CGRP + 1.0x olcegepant (p = 0.99). Diarrhea is a frequent observation after 1.0x CGRP and 1.0x CGRP + 1.0x sumatriptan but not after 1.0x CGRP + 1.0x olcegepant. Mice treated with SNP did not exhibit diarrhea or increased fecal granules across all time points or when compared to vehicle. In addition, piloerection, urination, and tremor did not significantly differ between CGRP or SNP-treated groups (data not shown). These results suggest that computing an index for motion sickness based on fecal incontinence, urination, piloerection, and tremor is problematic due to how biased the index is towards fecal excretion. Thus, we used a different method to assess motion-sickness based on motion-induced thermoregulation.

### Hypothermia and tail vasodilations occur during provocative motion and not stationary testing

**Fig. 2A** describes the experiment for assessing motion-induced thermoregulation. After a five-minute baseline recording, mice (n = 28M/27F) are challenged with provocative motion (t = 0 mins, rotation = ON) and exhibited gradual hypothermia until the end of the motion (t = 20 mins, rotation = OFF). 2^nd^ order curve fits calculated similar magnitudes of the mouse’s head hypothermia to between males and females (**Fig. 2B)**. In addition to the hypothermia, mice exhibited transient but significant increases in tail temperature during the first 10 minutes of the provocative motion. The Δ tail vasodilations were quantified for females (4.34 ± 0.34 °C) and males (5.44 ± 0.42 °C) (**Fig. 2D**). To ensure that the hypothermia and the tail vasodilations were physiological responses to the provocative motion stimulus, a separate group of mice (n = 10M/9F) was tested similarly but for stationary testing after vehicle treatment. No observed hypothermia or transient vasodilations were seen in mice of either sex during the stationary tests (**Fig. 2C** & **2E**).

### CGRP and SNP’s effects during stationary test

To exclude the possibility of thermoregulatory changes that may be due to CGRP and SNP effects, mice assessed for vehicle stationary tests (n = 9M/9F) were further tested with 1.0x CGRP **(Fig. 3A & D)** and 1.0x SNP **(Fig. 3B &** E). No differences were seen in the head or tail temperatures after 1.0x CGRP or 1.0x SNP testing compared to vehicle, and hypothermia or Δ tail vasodilations were not observed.

**Fig. 3:**
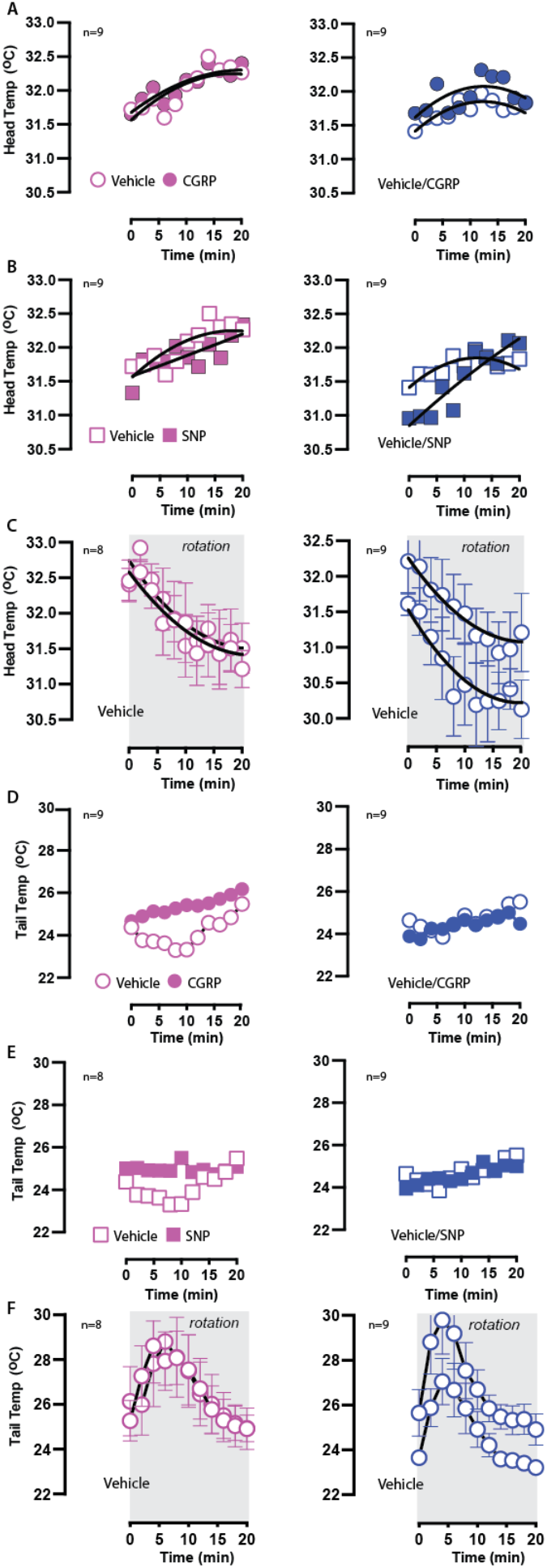
Stationary testing after IP CGRP/SNP and test-retest of IP vehicle control. Sample sizes are listed in the upper left corner. **(A and B)** Head temperature curves were recorded during stationary testing after IP delivery of vehicle, 1.0x CGRP, and 1.0x SNP. **(C and F)** A different cohort of mice was used to assess the test-retest reliability of head and tail temperatures during provocative rotation. **(C)** When assessing test-retest reliability, no significant differences were seen in the approximate magnitude of hypothermia in females (test vs retest: -1.2 °C vs -1.2 °C) or males (test vs retest: -1.3 vs 1.2 °C). (**D and E**) Tail temperatures were compared between stationary tests for vehicle, 1.0x CGRP, and 1.0x SNP as similarly done with head temperatures, and no significant differences were seen. **(F)** Similar to head temperatures, test-retest of tail measurements during provocative rotation showed no significant differences in Δ tail vasodilations in either female (test vs retest: 4.00 ± 0.80 °C vs 4.20 ± 0.80 °C) or male (test vs retest: 4.90 ± 0.80 °C vs 4.90 ± 0.90 °C).

### Test-retest reliability during provocative motion

After vehicle administration, male and female mice exhibited hypothermia during the provocative rotation and this response was repeatably observed in the retest (n = 9M/8F). In **Fig. 3C**, 2^nd^ order curve fits were generated for test-retest head curves and strong fits were seen for female (Pearson’s *r (23)* = 0.87, p < 0.0001) and for male (Pearson’s *r (23)* = 0.89, p< 0.0001). The magnitude of hypothermia measured in the head temperatures was similar between sexes and reproducible during the test-retest. In **Fig. 3F**, similar tail temperature profiles were observed during test-retest for females (Pearson’s *r* (23) = 0.92, p < 0.0001) and for male (Pearson’s *r (23)* = 0.95, p < 0.001). The Δ tail vasodilations were observed to have similar magnitudes for females (mean vasodilation _test_: 4.0 ± 0.8 °C, mean vasodilation _retest_: 4.2 ± 0.8 °C) and for males (mean vasodilation _test_: 4.9 ± 0.8 °C, mean vasodilation _retest_: 4.9 ± 0.9 °C).

### CGRP and SNP affect motion-induced thermoregulation in C57/B6J mice

As seen in **Fig. 4A and B**, 2^nd^ order curve fits were used to compute the recovery of head temperatures back to baseline values after the hypothermia after 1.0x CGRP delivery (28F/27M). A general trend of extended recovery times was observed due to 1.0x CGRP delivery (female: 24.0 mins, R^2^ = 0.73; male: 23.7 mins, R^2^ = 0.83) compared to their vehicle control responses (female: 20.7 mins, R^2^ = 0.76; male: 19.0 mins, R^2^ = 0.87). We observed no differences in the magnitude of head hypothermia between CGRP and vehicle control tests in either sex. **Fig. 4E and F** show tail temperature profiles after 1.0x CGRP, and a 2-way ANOVA showed effects of biological sex (F (1, 53) = 6.79, p = 0.01) and CGRP (F (1, 53) = 251.7, p < 0.001). 100% of mice exhibited normal Δ tail vasodilations after vehicle control administration but a majority of mice (89% female, 59% male) exhibited diminished Δ tail vasodilations (Bonferroni, p < 0.001) after 1.0x CGRP administration when compared to vehicle (**Fig. 5E**). Due to the diminished Δ vasodilations and the extended recovery times detected after CGRP testing, it is inferred that 1.0x CGRP impacted a mouse’s natural response to the provocative motion and their nausea response. In parallel, a different group of mice was used to assess SNP’s effects on motion-induced thermoregulation (8F/8M), and temperature profiles were analyzed. When measuring head temperatures, we did not observe any differences in recovery time or in the magnitude of head hypothermia (**Fig. 4C and D**). However, similar to CGRP, a 2-way ANOVA indicated SNP impacted Δtail vasodilations (F (1, 14) = 38.8, p < 0.001) but saw no effects of biological sex (F (1, 14) = 0.002, p = 0.97). Bonferroni post hoc analysis indicated 100% of females (p < 0.001) and 75% of males (p < 0.01) exhibited diminished vasodilations after 1.0x SNP (**Fig. 6G and 6H**).

**Fig. 4:**
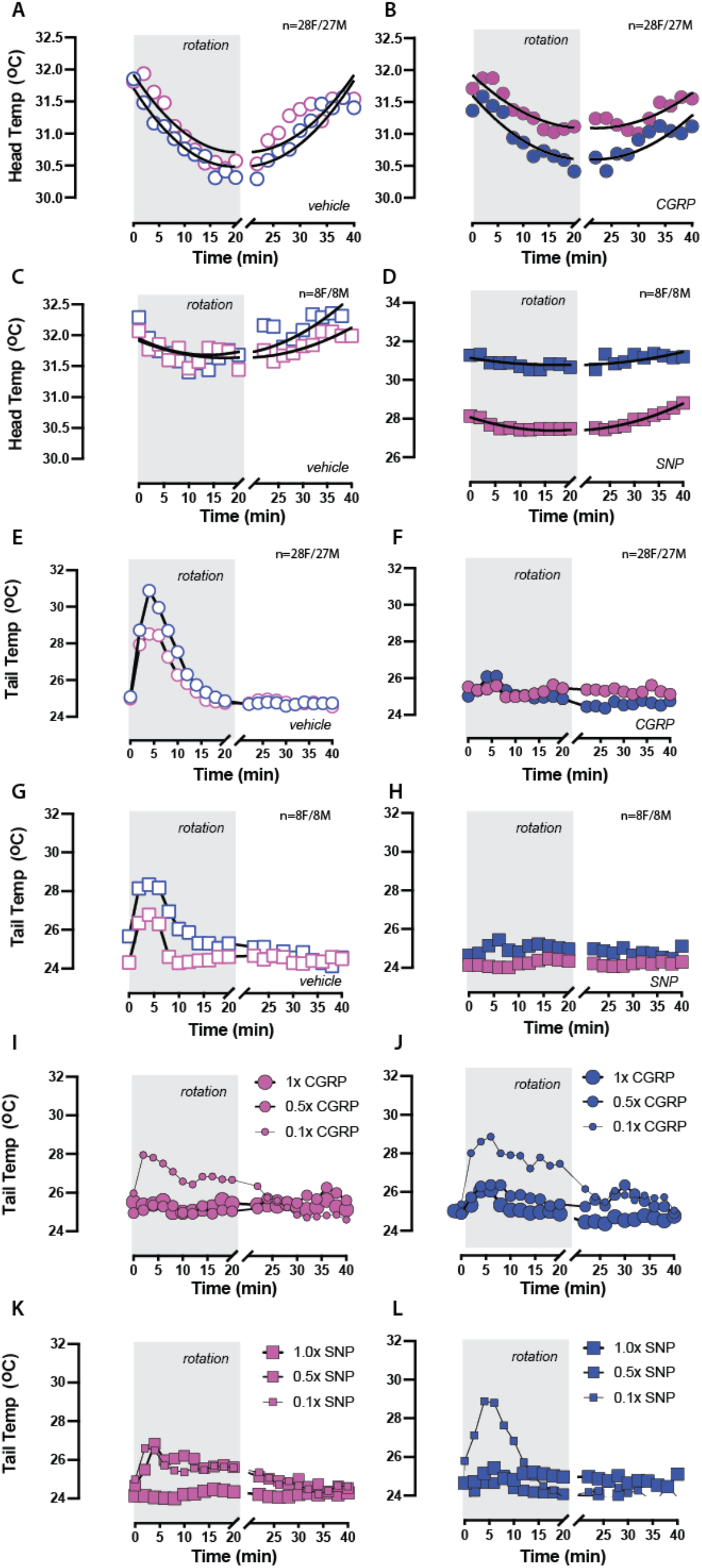
CGRP/SNP effects on hypothermia to provocative motion and dose-dependent changes in tail vasodilations. Sample sizes are shown in the upper right corner. Mice allocated for assessing CGRP’s effects on head and tail temperatures were tested for **(A & E)** vehicle and then **(B & F)** 1.0x CGRP. When assessing head temperature recovery, the 2^nd^-order curve fits extrapolated recovery times for vehicle control (female: 20.7 mins, male: 19.0 mins) and IP CGRP (females: 24.0 mins, male: 23.7 mins). The Δ tail vasodilations observed after vehicle treatment significantly diminished after 1.0x CGRP. A separate mice cohort was allocated for assessing SNP’s effects on head and tail and was tested for **(C & G)** vehicle control and later **(D & H)** 1.0x SNP. While head temperature recovery did not differ between vehicle and 1.0x SNP groups, Δ tail vasodilations were diminished after 1.0x SNP. **(I – L)** Dose-dependent changes in tail temperatures were observed for both IP CGRP and IP SNP. For CGRP testing, 0.1x, 0.5x, and 1.0x concentrations were prepared at 0.01, 0.05, and 0.1 mg/kg. For SNP testing, 0.1x, 0.5x, and 1x concentrations were prepared at 0.25 mg/kg, 1.25, and 2.5 mg/kg.

**Fig. 5:**
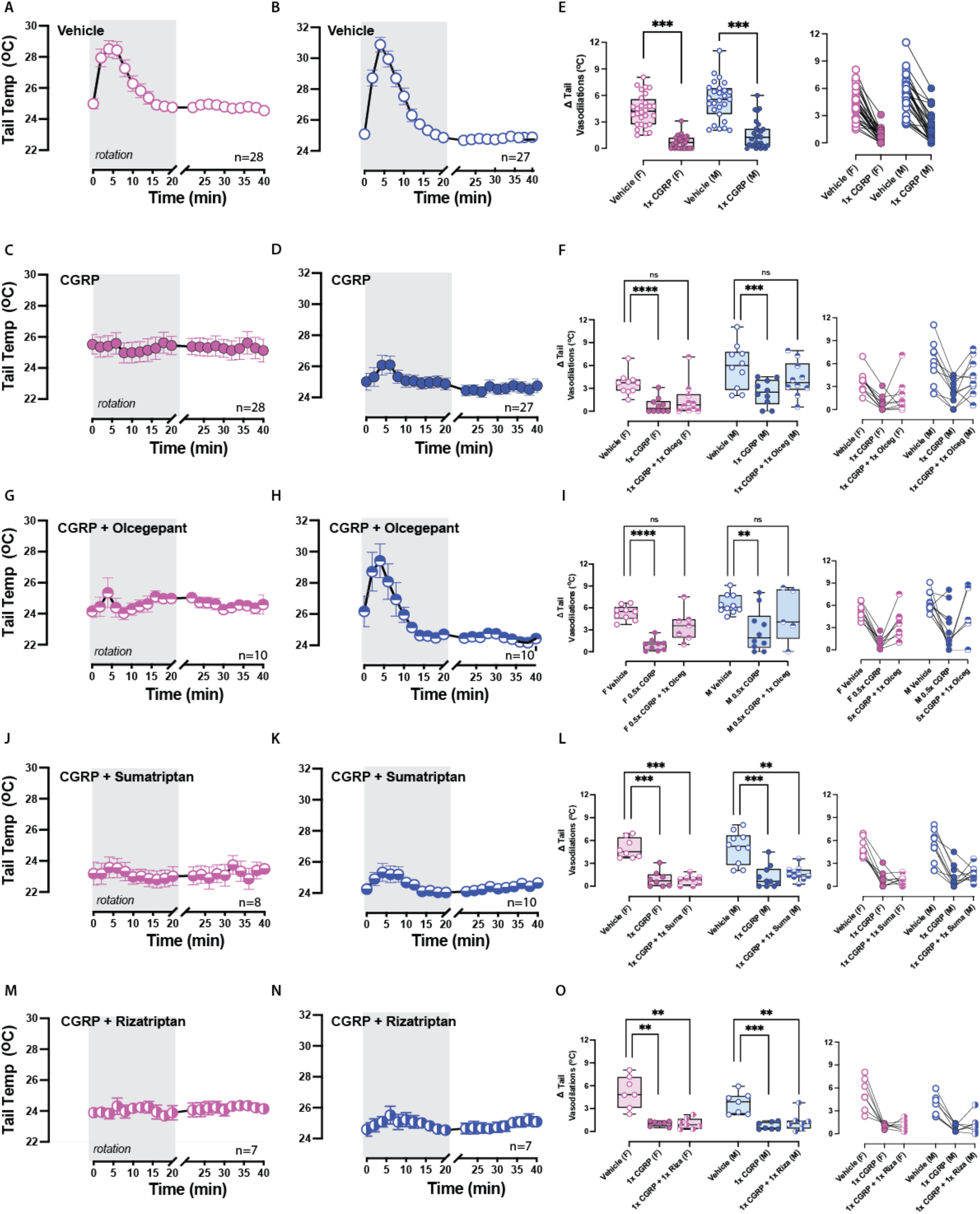
Olcegepant, but not sumatriptan or rizatriptan, protects against CGRP’s effects on tail vasodilations. Sample sizes are labeled in the bottom right corner. Tail temperatures were recorded after IP delivery of **(A & B)** vehicle, **(C & D)** 1.0x CGRP, **(F & G)** 1.0x CGRP + 1.0x olcegepant, **(J & K)** 1.0x CGRP + 1.0x sumatriptan and **(M & N)** 1.0x CGRP + 1.0x rizatriptan. **(E)** Bonferroni post-hoc indicated diminished tail vasodilations after 1.0x CGRP in all mice (p < 0.001 in either sex). **(F)** Dunnett post-hoc comparisons suggested olcegepant protected against CGRP-induced effects more strongly in males than females. When comparing 1.0x CGRP + 1.0x olcegepant to vehicle, no differences were observed in males. In females, differences were deemed not significant but were less certain as p = 0.057. **(I)** Mice treated with 0.5x CGRP and exhibited diminished Δ tail vasodilations were treated with 0.5x CGRP + 1.0x olcegepant and recovered their response. **(L & O)** Mice were not protected by 1.0x CGRP + 1.0x sumatriptan or by 1.0x CGRP + 1.0x rizatriptan as Δ tail vasodilations were still diminished compared to their vehicle response (vehicle vs CGRP + sumatriptan: p < 0.001 in females and p = 0.004 in males; vehicle vs CGRP + rizatriptan: p = 0.003 in females and p = 0.004 in males).

**Fig. 6:**
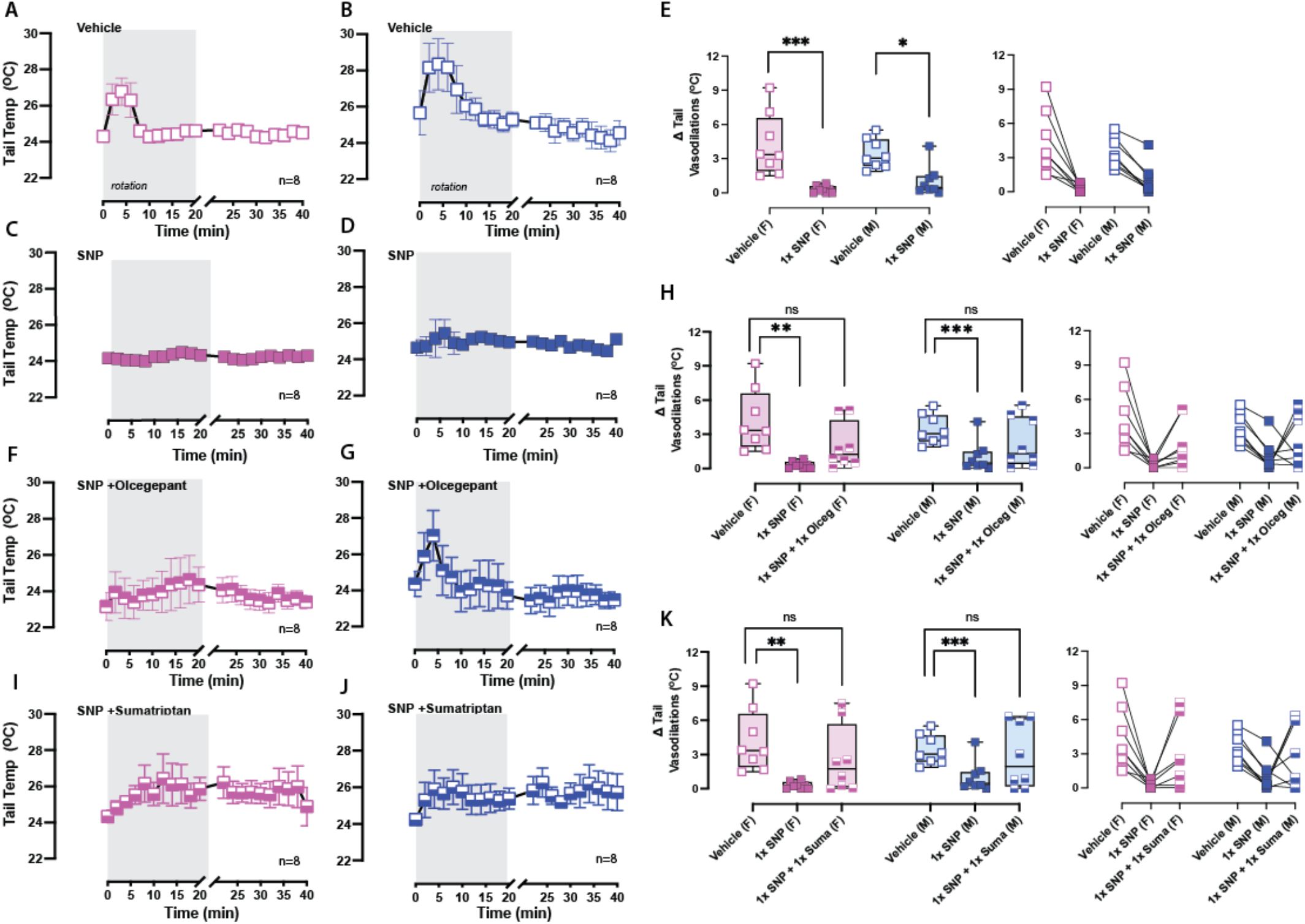
Olcegepant and sumatriptan block SNP’s effects on tail vasodilations: Similar to CGRP testing, tail temperatures were recorded after IP delivery of **(A & B)** vehicle, **(C & D)** 1.0x SNP, **(F & G)** 1.0x SNP + 1.0x olcegepant, and **(I & J)** 1.0x SNP + 1.0x sumatriptan, and sample sizes are labeled in the bottom right corner. **(E)** Dunnett’s post-hoc comparisons showed that SNP diminished Δ tail vasodilations compared to vehicle control in both sexes. No differences were observed between **(H)** vehicle vs 1.0x SNP + 1.0x olcegepant, or **(K)** vehicle vs 1.0x SNP + 1.0x sumatriptan in either sex.

### Dose-dependent response of CGRP or SNP on tail vasodilations

While 1.0x concentrations were used for the motion sickness assay in this study and these concentrations are based on previous studies assessing these migraine triggers, it was unclear whether lower doses would also elicit observed responses. To answer this question, a dose-dependent response curve was tested in mice (n = 10M/9F) and Δ tail vasodilations were analyzed for vehicle, 0.1x and 0.5x of IP CGRP or IP SNP, and these mice were compared with the original groups of mice tested for vehicle and 1.0x CGRP (**Fig. 4I-L**). A 2-way mixed effects model and Bonferroni post hoc assessed the factors: biological sex and CGRP/SNP dilutions. 0.1x CGRP had no significant effect on Δ tail vasodilations in either sex. However, differences were observed when comparing Δ tail vasodilations between vehicle and 0.5x CGRP **(Supplementary Fig.S1 A.)** in females (p < 0.001) and in males (p = 0.03). Significant differences in Δ tail vasodilations were seen when comparing vehicle and 1.0x CGRP in both sexes (p < 0.001 each). When assessing SNP’s effects (n = 8F/8M), no significant differences were seen at 0.1x and 0.5x SNP compared to vehicle control **(Supplementary Fig. S1 B)**, but a significant difference was seen when comparing vehicle to 1.0x SNP in females (p = 0.01) and in males (p = 0.0002). The greatest effect of CGRP and SNP on Δ tail vasodilations was observed at their 1.0x concentration and these concentrations were used to assess the migraine blockers.

### Olcegepant but not triptans protect against CGRP’s effects on tail vasodilations

Administration of 1.0x CGRP reduced Δ tail vasodilations in nearly all mice (**Fig. 5E**), and so it was hypothesized that these mice would regain their healthy response to provocative motion with IP treatment of either CGRP + olcegepant (n = 10M/10F), CGRP + sumatriptan (m = 10M/8F), or CGRP + rizatriptan (n = 7M/7F), and different groups of mice were used. Prior to assessing CGRP + blockers, groups of mice were assessed for blocker-only effects by testing vehicle + 1.0x olcegepant (n = 8F/10M), vehicle + 1.0x sumatriptan (n = 7F/10M), or vehicle + 1.0x rizatriptan (n = 7F/7M), but no differences were seen when compared to their vehicle control so the blockers, by themselves, do not have a secondary effect on tail vasodilations **(Supplementary Fig. S2)**.

Tail temperature profiles were recorded and displayed for vehicle and drug treatments (**Fig. 5A-D, G, H, J, K, M & N**). After quantifying Δ tail vasodilations, one-way ANOVAs were used to assess CGRP + blocker effects in the designated groups, and Dunnett post-hoc analysis was used to compare treatments to vehicle. F-values and ANOVA p-values are listed in **Table 1**. We observed olcegepant blocked CGRP-induced changes in males because Δ tail vasodilations resembled vehicle **(Fig. 5F)**, and this protective effect was trending in females (p = 0.06). In contrast, triptans did not have a protective effect **(Fig. 5L & O)**. Mice after 1.0x CGRP + 1.0x sumatriptan still exhibited diminished Δ tail vasodilations compared to their IP vehicle test (p < 0.001 in females and p = 0.004 in males). A similar observation was seen in mice after treatment of IP 1.0x CGRP + 1.0x rizatriptan, as tail vasodilations were still significantly diminished in both sexes compared their IP vehicle test (p = 0.003 in females and p = 0.004 in males). In the dose-dependent response study, mice that experienced reduced vasodilations due to the 0.5x CGRP dose were later tested with 0.5x CGRP + 1.0x olcegepant, and 86% of females and 80% of males were rescued after delivery of 1.0x olcegepant (**Fig. 5I)**. These findings show that olcegepant blocked the effects of CGRP and allowed for a majority of mice to exhibit their normal response to the provocative motion by blocking CGRP’s effects, while both triptans were not effective.

### SNP’s effects on tail vasodilations are blocked by olcegepant

The effects of olcegepant and sumatriptan were also assessed against 1.0x SNP (n = 8M/8F) that had reduced Δ tail vasodilations in nearly all mice **(Fig. 6E)**, and tail temperatures are depicted (**Fig. 6A-D, F, G, I, & J)**. In order to conserve mice, the same mice were treated with 1.0x CGRP + 1.0x olcegepant and four days later with 1.0x CGRP + 1.0x sumatriptan. 1-way ANOVAs with Dunnett post hoc were computed and F-values and statistics are in **Table 1**. In females and males, robust Δ tail vasodilation responses were observed in 50% of females and 50% males after 1.0x CGRP + 1.0x olcegepant **(Fig. 6H)**, and these percentages were also seen when mice were later tested with 1.0x CGRP + 1.0x sumatriptan **(Fig. 6K)**. Post-hoc analyses in both sexes indicated no significant difference in tail vasodilations for either SNP + blocker when compared to vehicle. These mice were not further tested with blockers at lower doses of SNP. **Table 2** is provided to depict the percentage differences in the prevalence of diminished Δ tail vasodilations observed across all treatments (CGRP/SNP + blockers) for both male and female mice.

**Table 2:** Diminished tail vasodilations indicate a disrupted response to provocative rotation. Percentage of mice that exhibited diminished Δtail vasodilations (<1.5° C) during motion-induced thermoregulation testing across treatment. Treatments are administered at the following concentrations: 0.1x CGRP - 0.01 mg/kg, 0.5x CGRP 0.05 mg/kg, 1.0x CGRP - 0.1 mg/kg, 1.0x olcegepant - 1 mg/kg, 1.0x sumatriptan 0.6 mg/kg, 1.0x rizatriptan - 0.6 mg/kg, 1.0x SNP - 2.5 mg/kg.

| **Reduced $\Delta$ tail vasodilations (< 1.5 °C)**<br>**during motion-induced thermoregulation** |  |  |
| --- | --- | --- |
| **Treatment** | **Female %** | **Males %** |
| Vehicle | 0.0% | 0.0% |
| 0.1x CGRP | 22.2% | 0.0% |
| 0.5x CGRP | 77.7% | 50.0% |
| 1.0x CGRP | 89% | 59.0% |
| 0.5x CGRP + 1.0x olcegepant | 14.2% | 20.0% |
| 1.0x CGRP + 1.0x olcegepant | 70.0% | 10.0% |
| 1.0x CGRP + 1.0x sumatriptan | 77.8% | 50.0% |
| 1.0x CGRP + 1.0x rizatriptan | 71.4% | 85.7% |
| 0.1x SNP | 20.0% | 0.0% |
| 0.5x SNP | 50.0% | 50.0% |
| 1.0x SNP | 100% | 75.0% |
| 1.0x SNP + 1.0x olcegepant | 50.0% | 50.0% |
| 1.0x SNP + 1.0x sumatriptan | 50.0% | 50.0% |

## Discussion

In this study, we show that systemic CGRP injections can induce motion sickness in wildtype C57B6/J mice, and this motion sickness can be strongly antagonized by the CGRP receptor blocker, olcegepant. Published studies assessing CGRP’s effects on light-aversive behavior (photophobia) and cutaneous hypersensitivity (allodynia) in rodents indicate triptans are effective in reducing these surrogate behaviors ^30, 31^, but interestingly in our study, sumatriptan and rizatriptan did not show significant efficacy in blocking CGRP’s effects in motion sickness and motion-induced nausea. We have used two different murine motion sickness assays: 1) MSI and 2) motion-induced thermoregulation. In both assays, provocative motion caused distinct changes, but MSI was too weighted on gastric distress. This bias reduced its validity as an assay because gastrointestinal complications do not directly imply motion sickness. However, the motion-induced thermoregulation model is more suitable for studying motion sensitivity as it involves studying motion-induced thermoregulation caused specifically by a vestibular stimulus – the constant orbital rotation. In addition, the motion-induced thermoregulation model was robust for assessing CGRP and SNP’s effects. Similar rodent studies suggest tail vasodilations are a precursor to emesis and removal of toxins ^11^. Our data showing CGRP’s blunting of tail vasodilations to provocative motion suggests that motion-induced nausea may be aggravated with CGRP and that olcegepant is protective but not triptans. This result is in agreement with clinical findings that showed a 2-hour infusion of CGRP caused gastrointestinal hyperactivity and nausea, and pretreatment with a CGRP receptor antagonist ameliorated these clinical symptoms but pretreatment with sumatriptan did not ^32^. Moreover, a clinical meta-analysis of anti-CGRP signaling treatments for nausea in episodic migraine provided strong support for their effectiveness ^33^. Interestingly, we also observed that female mice were more severely affected by systemic CGRP than their male counterparts, which correlates with preclinical studies showing stronger tactile sensitivity in female mice when challenged with durally applied CGRP ^34^.

### CGRP signaling and murine motion-sickness indices

Others have shown in human subjects and in mouse models that CGRP causes diarrhea by disrupting peristaltic intestinal activity and promoting ion and water secretion in the intestinal lumen, and that CGRP signaling antagonism can reverse these effects ^35–37^. Thus, it is expected that CGRP-induced diarrhea would cause noticeable increases in the motion sickness index (MSI) calculation. The OVAR rotation further increased the motion sickness index (MSI) regardless of CGRP or SNP, highlighting the effect of a strong vestibular perturbation in increasing MSI outcomes.

Interestingly, the motion-induced thermoregulation model revealed differences between vehicle and CGRP/SNP injected mice. Differences in motion-induced nausea after CGRP administration were blocked by a CGRP receptor antagonist (olcegepant) but not by triptans (sumatriptan or rizatriptan). We also showed that in the stationary tests, there are no thermoregulatory changes in the absence of provocative motion. While male and female mice both showed disrupted thermoregulatory responses to CGRP injections, dose response experiments at 0.05 and 1 mg/kg CGRP show that olcegepant can rescue CGRP-induced nausea in male yet olcegepant did not fully rescue all female mice. (**Table 2**). This sexual dimorphism is similar to higher incidence of migraine and VM in female patients ^4, 38, 39^ and reports of sexual-dimorphic effects of CGRP both in dura-induced pain, spinal cord neuropathic pain, and preclinical surrogate of allodynia or touch sensitivities ^34^. To this end, other studies have demonstrated a sex difference in the expression of CGRP receptor components in the trigeminal nucleus ^40^.

This study is the first to provide preclinical evidence for the use of CGRP antagonists in treating motion hypersensitivities that occur in VM, and should be paired with findings from pertinent clinical trials for validation. We provide data that the motion-induced thermoregulation assay is sensitive to the known migraine triggers, CGRP and SNP, and show the effectiveness of different migraine drugs (‘gepants’ and triptans) in reducing nausea responses observed in this mouse model. Future experiments will assess if mice over- or under-expressing the CGRP receptor will be more or less susceptible to changes in their thermoregulatory responses to provocative motion, and if migraine blockers can provide protection. These experiments will achieve the greater goal of elucidating the role of CGRP signaling in the symptoms of nausea and motion sickness that occur in migraine and VM.

## Study Highlight Bullet points

- Enhanced motion-induced nausea is an observed symptom in migraine and especially vestibular migraine (VM).
- We used two different murine motion sickness assays (thermoregulation and MSI), but MSI was too weighted on gastric distress to be effective.
- Differences in motion-induced thermoregulation after systemic CGRP were blocked by olcegepant but not by triptans.
- Olcegepant was more effective in rescuing CGRP-induced thermoregulatory changes in male compared to female mice.

## Acknowledgments

We would like to acknowledge assistance with data collection and analysis from Vedat Duzgezen, Stefanie Faucher, Elana Fine, Catherine Hauser, Benjamin Liang, and Blaze Strangio. This work was fully supported by NIH R01DC017261 (AEL).

---

**Supplementary Fig. S1:**
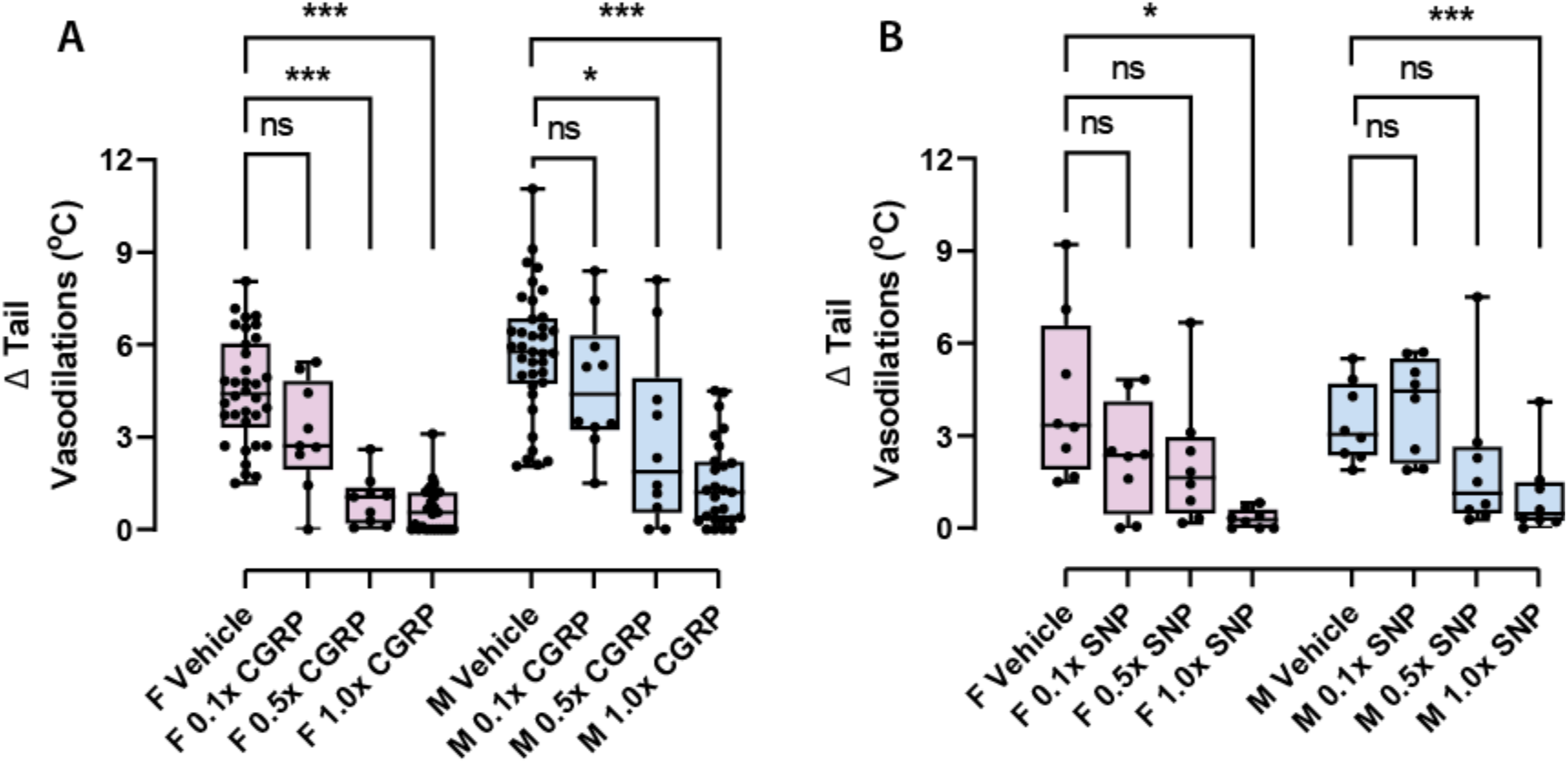
**A.** 2-way mixed effects model assessed the factors i) sex (male vs female) and ii) CGRP dilutions (0.1x, 0.5x, and 1.0x CGRP). Compared to vehicle responses, significant differences were observed at 0.5x CGRP for females (p < 0.001) and for males (p = 0.03). These differences were more strongly observed at 1.0x CGRP (p < 0.001 for both sexes). 0.1x CGRP did not have a strong effect. Sample sizes are listed: vehicle (34 F/37M), 0.1x, and 0.5x CGRP (9F/10M), 1.0x CGRP (28F/27M). **B**. 2-way mixed effects model assessed the factors i) sex (male vs female) and ii) SNP dilutions (0.1x, 0.5x, and 1.0x SNP). Compared to vehicle responses, significant differences were observed at 1.0x SNP for females (p = 0.015) and for males (p = 0.0002). Significant differences were not seen at 0.1x and 0.5x SNP. All 8F/8M were tested across dilutions.

**Supplementary Fig. S2:**
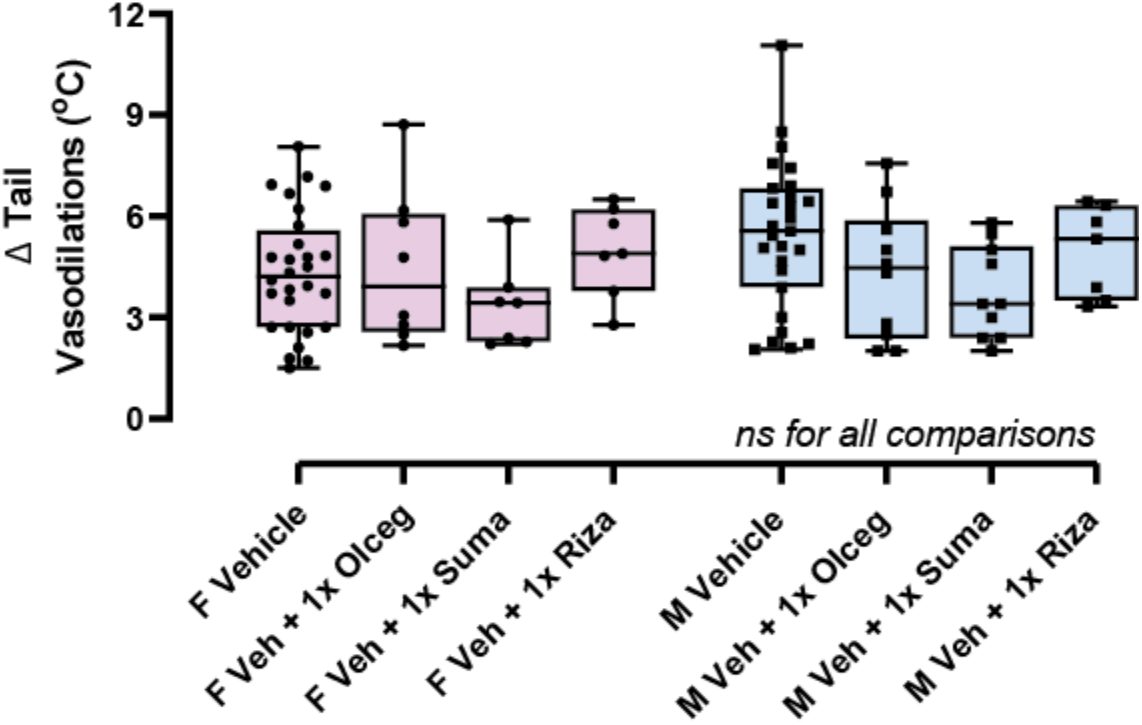
In females and males, 1-way mixed effect models with Dunnett’s post hoc comparison tests were computed to assess drug controls to the vehicle (Veh), and no significant differences were observed. Sample sizes are listed: vehicle (28 F/27M), Veh + 1x Olceg (8F/10M), Veh + 1.0x Suma (7F/10M), Veh + 1.0x Riza (7F/7M).

